# The *Chlamydia trachomatis* secreted effector protein CT181 binds to Mcl-1 to prolong neutrophil survival

**DOI:** 10.1101/2025.03.16.643443

**Authors:** Robert Faris, Rebecca Koch, Paige McCaslin, Naveen Challagundla, Brianna Steiert, Shelby E. Andersen, Parker Smith, C.A. Jabeena, Peter Yau, Thomas Rudel, Mary M. Weber

**Affiliations:** Department of Microbiology and Immunology, University of Iowa Carver College of Medicine, Iowa City, Iowa, USA; Department of Microbiology, Biocenter, University of Wuerzburg, Wuerzburg, Germany Chair of Microbiology, University of Würzburg, Würzburg, Germany; Department of Veterinary Microbiology and Pathology, Washington State University, Pullman WA, USA; Department of Immunology and Microbiology, University of Colorado - Anschutz Medical Campus, Aurora, CO, USA; Emory National Primate Research Center, Emory University, Atlanta, GA, USA; Carver Biotechnology Center – Protein Sciences Facility, University of Illinois at Urbana–Champaign, Urbana, IL, USA

**Author notes:** RF, RK, and PM contributed equally to this work.

**Keywords:** CT181, chlamydia, T3SS effector, neutrophil, Mcl-1

## Abstract

*Chlamydia trachomatis* (*C.t*) infections can lead to severe complications due to the pathogen’s ability to evade the host immune response, often resulting in asymptomatic infections. The mechanisms underlying this immune subversion remain incompletely understood but likely involve specific bacterial effector proteins. Here, we identify CT181 as a novel effector that directly binds to Mcl-1, a key regulator of neutrophil survival. While a *C.t.* CT181 mutant exhibited only modest defects in epithelial cell replication and inclusion development, it was essential for *C.t.* survival in neutrophils, correlating with Mcl-1 stabilization. Using a murine infection model, we demonstrate that CT181 is required for *C.t.* colonization and cytokine production *in vivo*. Our findings establish CT181 as the first bacterial effector protein known to bind Mcl-1 to enhance neutrophil survival, revealing a critical strategy by which *C.t.* promotes immune dysregulation, facilitating bacterial persistence while driving *C.t.* pathogenesis.

## INTRODUCTION

The obligate intracellular pathogen *Chlamydia trachomatis* (*C.t.*) is the causative agent of trachoma (1) and the sexually transmitted infection chlamydia (2). While many cases are asymptomatic, untreated infections can result in serious complications, including pelvic inflammatory disease, ectopic pregnancy, sterility, and the development of cervical or ovarian cancer (2–5). There is no vaccine and over 127 million new cases are reported annually worldwide (6). Moreover, antibiotic failure occurs in 10% of cases (7, 8).

All *Chlamydiae* share a unique biphasic developmental cycle in which they differentiate between infectious elementary bodies (EB) and replicative reticulate bodies (RB) (9). Upon contact with a target host cell, EBs are internalized into a membrane-bound inclusion (9), which avoids lysosomal fusion and traffics along microtubules to the peri-Golgi region (10, 11). EBs differentiate into RBs, undergoing multiple rounds of replication prior to converting back to EBs for release by host cell lysis or extrusion (12). At all stages of the developmental cycle, *C.t.* utilizes its type III secretion (T3S) system to secrete effector proteins to engage host organelles and perturb vital signaling pathways to acquire key nutrients for replication, to promote host cell viability, and to subvert host defense mechanisms.

As an obligate intracellular pathogen, *C.t.* is dependent on the host cell to provide a replicative niche conducive to completing its replication cycle. To achieve this, *C.t.* has evolved methods to inhibit the induction of apoptosis and to promote host cell viability. *C.t.* employs multiple strategies to subvert host cell death, including inhibiting activation of Bax and Bak, preventing mitochondrial outer membrane permeabilization and cytochrome c release, and inhibiting caspase-3 cleavage (13, 14). Additionally, pro-survival signaling pathways, including Raf/MEK/ERK and PI3K/AKT, are activated during *C.t.* infection, which in turns induces the up-regulation and stabilization of induced myeloid leukemia cell differentiation protein (Mcl-1) (15). The importance of this process is underscored by studies indicating that the depletion of Mcl-1 sensitizes the *C.t.-*infected cell to death (15). Prior work revealed that Cdu1 localizes to the inclusion membrane and stabilizes Mcl-1 through deubiquitination (16). However, infection with a Cdu1 mutant only resulted in a marginal decline in Mcl-1 protein levels (16), suggesting that Mcl-1 levels are regulated via multiple mechanisms and might involve additional *C.t.* effector proteins.

While the identification of inclusion membrane proteins (Incs), has readily been achieved by screening proteins for the presence of a bi-lobed hydrophobic domain, the identification of other type III secreted proteins has proven more challenging. Although some have been identified based on the presence of a eukaryotic like domain, signal sequence, and analysis in a surrogate host (17–20) it is likely that other undiscovered secretion substrates remain to be identified. *C.t.* effectors TmeA, TarP, TmeB, and TepP are uniquely produced by the EB and have been shown to be secreted early in infection to facilitate host cell invasion and initiate early infection events (21–24). Thus, we hypothesized that other hypothetical proteins, uniquely present in EBs, may represent novel secreted effector proteins (25). Here we assessed whether any of these hypothetical proteins uniquely produced in EBs (25) are secreted during *C.t.* infection, successfully identifying three novel secretion substrates. Using yeast-two-hybrids and affinity purification mass spectrometry, we determined that the secreted effector protein CT181 binds to the induced myeloid leukemia cell differentiation protein (Mcl-1). Importantly, we show that the CT181 mutant was severely impacted in its ability to grow in neutrophils and in its ability to survive in the mouse genital track. By binding to Mcl-1, CT181 serves to promote chlamydial disease by impacting host cell survival pathways, highlighting its crucial role in both immune evasion and pathogen persistence during infection.

## MATERIALS AND METHODS

### Bacterial and cell culture

*Chlamydia trachomatis* serovar L2 (LGV 434/Bu) was propagated in HeLa 229 cells (American Type Culture Collection) and EBs were purified using a gastrografin density gradient as previously described (26). HeLa cells were grown at 37°C with 5% CO_2_ in RPMI 1640 medium (ThermoFisher Scientific) supplemented with 10% Fetal Bovine Serum (Gibco). A2EN cells (Kerafast) were propagated in keratinocyte-serum free media (K-SFM) (ThermoFisher Scientific) supplemented with 0.16 ng/ml epidermal growth factor (EGF), 25 μg/ml bovine pituitary extract (BPE), 0.4 mM CaCl_2_, and gentamicin (27).

### Human and mouse PMN isolation

Mouse neutrophils were isolated from the bone marrow of naïve C57BL/6N mice by density gradient centrifugation. Bone marrow cells were overlaid on to Histopaque 1119 and Histopaque 1077 and centrifuged for 30 min at 872 RCF at room temperature without brake. Neutrophils were collected at the interface of Histopaque 1119 and Histopaque 1077, washed with RMPI 1640, and used immediately. Human neutrophils were freshly isolated from the blood of healthy volunteers using the Ficoll separation method (44). Briefly, blood was layered on top of Ficoll and centrifuged at 1500 RCF for 30 min. All cell layers above the PMN layer were removed and Polyvinyl alcohol (PVA) solution was added to the remaining layers. After incubation for 45 min the light red layer was transferred to a fresh tube and centrifuged at 1000 RCF for 5 min. The remaining red blood cells were lysed with sterile H_2_O and the osmolarity was then restored by adding sterile 5X PBS. These neutrophils were collected by centrifugation and then resuspended in RPMI medium. The neutrophil number was determined using a Neubauer chamber and trypan blue staining.

### Plasmid construction

To assess secretion of candidate effectors, each candidate secretion substrate was PCR amplified from L2/434/Bu genomic DNA and cloned into the NotI/KpnI site of pBomb4 CyaA, pBomb4 BlaM, and pBomb4 GSK FLAG (28). For affinity-purification mass spectrometry, CT181 was cloned into the NotI/KpnI site of pBomb4-tet-mCherry (29) and a FLAG tag was added to the C-terminus via PCR. TargeTron insertion sites were predicted by TargeTronics and gene blocks were obtained from Integrated DNA Technologies. Gene blocks were cloned into the HindIII/BsrGI site of pACT to generate CT181*::bla* (30). For ectopic expression, CT181 was cloned into the KpnI/XhoI of pcDNA-GFP. The integrity of all constructs was verified by DNA sequencing at McLab. All primers are listed in Table S1.

### Transformation of *Chlamydia*

*C.t.* serovar L2 (LGV 434/Bu) EBs were transformed as previously described (28). Infectious progenies were harvested every 48 h and used to infect a new HeLa cell monolayer until viable inclusions were evident (∼2-3 passages). Expression of individual fusion proteins was confirmed by western blotting. For the TargeTron mutant, successful insertion into the target gene, CT181, was confirmed by PCR.

### Adenylate Cyclase (CyaA) secretion assay

Confluent HeLa cell monolayers were infected at an MOI of 5 with *C.t.* transformant strains harboring the pBomb4 CyaA expression plasmids. Expression of the CyaA fusion protein was induced using 10 ng/ml anhydrous tetracycline (aTc) at the time of infection, and 24 h post-infection, the relative abundance of cAMP in host cells was measured via competitive ELISA as previously described (28). The levels of cAMP in cells infected with *C.t.* pBomb4 CyaA (negative control vector) was compared to those infected with *C.t.* CyaA-effector fusions strains to evaluate effector secretion.

### Beta-lactamase assay

To assay for effector secretion, HeLa cells, seeded into black, clear bottom 96-well plates (Greiner) were infected at an MOI of 5 and effector expression was induced at time of infection using 10 ng/ml aTc as previously described (28). At 24 h post-infection, cells were loaded with CCF4-AM using the alternative loading protocol following the manufacturer’s instructions (ThermoFisher Scientific). Plates were incubated in the dark for 1 h at room temperature and then were read on a plate reader (Tecan). To quantify effector translocation, the background was subtracted, the ratio of 460 nm to 535 nm (blue:green) was determined, and expression, relative to cells infected with *C.t.* expressing BlaM only, was calculated as previously described (31).

### GSK-FLAG immunoprecipitation

To evaluate effector secretion using the GSK assay, HeLa cells were infected at an MOI of 5 and effector-GSK FLAG fusion protein expression was induced using 10 ng/ml aTc as previously described (28). Cells were harvested 24h post-infection by lysing in 800 µl eukaryotic lysis solution (ELS) (50mM Tris-HCl, 150 mM sodium chloride, 1 mM ethylenediaminetetraacetic acid, and 1% Triton-X 100) containing Halt cocktail protease and phosphatase inhibitor (ThermoFisher Scientific) along with 10 µM GSK-3 α and β inhibitor 1-(7-Methoxyquinolin-4-yl)-3-[6-(trifluoromethyl)pyridin-2-yl]urea (Tocris). Supernatants were applied to anti-FLAG magnetic beads (ThermoFisher Scientific) for 1 h at 4°C and unbounded proteins were removed by washing the beads 5 times in ELS without Triton-X 100. Purified protein was eluted using 4X LDS sample buffer (ThermoFisher Scientific) and samples were analyzed by western blotting.

### Western blotting

To evaluate expression, confluent HeLa cell monolayers were infected at an MOI of 5 and after 24 h the samples were lysed in ELS with Halt cocktail protease inhibitor. Lysates were resolved using 3-8% Tris-Acetate protein gels with Tris-Acetate SDS running buffer for CyaA and BlaM fusion proteins. GSK FLAG IPs were resolved using 4-12% Bis-Tris protein gels with MOPS SDS running buffer. Proteins were transferred to a PVDF membrane and probed using anti-CyaA (Santa Cruz Cat# sc-13582), anti-BlaM (QED BioScience Cat# 15720), GSK-3β-Tag (Cell Signaling Cat# 9325S), or Phospho-GSK-3-beta (Cell Signaling Cat# 9336S) antibodies. For IPs, blots were probed with anti-FLAG (Thermo Cat# 701629) and anti-HA (Sigma Cat# H6908-100ul) antibodies.

To evaluate the expression of proteins in neutrophils, human or mouse neutrophils were infected at an MOI of 5 and after 48 h the samples were lysed either in 8M Urea or 3M trichloroacetic acid to precipitate the protein. The lysate was mixed with 2X Laemmli buffer and resolved using 10% acrylamide gel with Tris-Chloride SDS running buffer. Proteins were transferred to a PVDF membrane in Transblot and probed with anti-Mcl-1 (Cell Signaling Cat# 5453T), *C.t.* OmpA (Thermo Cat# PA5-117609) or anti-β-actin (Thermo Cat# A00702-100) antibodies.

### Immunofluorescence

To determine the subcellular localization of CT181 and co-localization with Mcl-1, HeLa cells were transfected using Lipofectamine LTX (ThermoFisher Scientific). Eighteen hours post-transfection, cells were fixed with 4% formaldehyde, permeabilized with 0.1% Triton-X 100, and the nucleus was stained using DAPI and mitochondria were stained using CoxIV (Novus MAB6980). Images were captured using a Leica DFC7000T confocal microscope equipped with Leica software.

### Growth Curve Assay

A2EN or HeLa cells were infected on ice at a MOI of 5 for A2EN or 2.5 for HeLa with each strain. After 30 min on ice, the inoculum was removed, and cultures were shifted to 37°C to stimulate bacterial uptake. At 0, 24, and 48h post-infection, cells were lysed in water and the supernatants were added to fresh HeLa cell monolayers as previously described (32). Titer plates were fixed 24h post-infection and stained with an anti-*C.t.* LPS antibody (Novus Cat#NBP1-28820) for enumeration of inclusions by immunofluorescence microscopy. In parallel, a set of samples was fixed at 24 h and stained with anti-*C.t.* LPS and anti-IncE (inclusion membrane marker). Inclusion area was measured using ImageJ.

Freshly isolated human or mouse PMNs were infected at an MOI of 5. At 48, 72 and 96 h post-infection, cells were sonicated at 4°C for 15 min and the supernatants were added to fresh overnight grown Hela cells (in case of human PMNs) and McCoy cells (in case of mouse PMNs) in 24 well plates. Cells were fixed 36 h post-infection and stained with an anti-*C.t.* HSP60 antibody (Santa Cruz Cat#SC-57840) and DAPI for enumeration of inclusions by immunofluorescence microscopy. Infection was calculated from the number of cells and inclusions present in randomly captured microscopic images.

### Invasion assay

To determine if CT181 is important for host cell invasion, A2EN cells were seeded on glass coverslips and an invasion assay was conducted as previously described (22). Briefly, cells were infected on ice at a MOI of 5 for 30 min after which the inoculum was removed and plates were placed at 37°C with 5% CO_2_ for 60 min to allow for bacterial uptake. Cells were fixed with 4% formaldehyde and differential immunostaining was conducted as previously described (33). The number of internal bacteria (single stained) and number of host cells (DAPI stained) was enumerated from at least 30 images per experiment.

### Yeast-two hybrid

ULTImate yeast two-hybrid analysis was completed by Hybrigenics Services (Paris, France). The coding sequence for CT181 *C.t.* L2/434/Bu (aa1-236) was PCR amplified and cloned into the pB27 construct as a C-terminal fusion with LexA (N-CT181-LexA). The resulting construct was introduced into yeast as bait and screened by mating with yeast bearing a randomly primed HeLa cell cDNA library (prey). Due to autoactivation, the screen was conducted on selective media with 100 mM-AT. Positively selected clones were isolated and identified using the NCBI GenBank. The predicted biological score (PBS) was calculated to assess the reliability of each interaction. Scores ranged from high probability of specificity (A score) to low probability of specificity (E score) between the bait and prey.

### Affinity purification mass-spectrometry

A confluent monolayer of HeLa cells was infected at an MOI of 2 with *C.t.* expressing FLAG-tagged CT181. After 24 h, cells were lysed in ELS, placed on ice for 20 min, and subsequently spun at 12,000 x g for 20 min. Supernatants were incubated with preclearing beads (mouse IgG agarose, Millipore Sigma) for 2 h and then were applied to FLAG magnetic beads (anti-FLAG M2 Affinity Gel, Millipore Sigma) overnight at 4°C. Beads were washed 6 times with ELS without Triton-X 100 in MS grade water and prepared for mass spectrometry as previously described (34, 35). Raw LC-MS/MS data were searched against a database containing UniProt_Human and Chlamydia_trachomatis_D/UW-3/CX using Mascot 2.8.

### Immunoprecipitation

HeLa cells were co-transfected with pcDNA3.1-GFP plasmids containing empty vector, CT181, or TmeA and pcDNA3.1-Mcl-1HA using Lipofectamine LTX (Thermo Fisher Scientific). Four h post transfection, the media was changed. For infection IPs, HeLa cells were infected at an MOI of 2 and the effector-FLAG fusion was induced using 10 ng/ml aTc as previously described(28). At 24 hpi, cells were washed with ice-cold 1X PBS, and lysed in 800 µl ELS containing Halt cocktail protease and phosphatase inhibitor(28). Supernatants were applied to anti-FLAG (infection) or anti-HA (transfection) magnetic beads (Pierce^TM^ Thermo Fisher Scientific) for 1.5h at 4°C. The beads were subsequently washed 5X in ELS without Triton-X 100 and the purified protein was eluted using 4X NuPAGE LDS Sample Buffer (ThermoFisher Scientific). Samples were analyzed by western blotting.

### Cell death assay

To determine neutrophil viability during *C.t.* infection, freshly isolated 10^6^ human or mouse neutrophils were infected with either wild-type *C.t.* or CT181::*bla* at an MOI 5 in 6 well plates in RPMI 1640 medium. After different time points, Annexin V-APC and 7-AAD (BD Pharmingen Cat#550475) were added to the cells in binding buffer and incubated for 15 min at 37°C. Cells were washed and analyzed using an Attune NXT flow cytometer. Cells negative for both Annexin V and 7-AAD were considered as viable cells and plotted as percent of double negative cells.

### *In vivo* mouse infection

C57BL/6N mice were injected with 2.5 mg depot medroxyprogesterone acetate (DMPA) 5 d prior to infection. On the day of the infection, mice housed in specific pathogen-free conditions, were trans-cervically infected with 10^7^ inclusion forming units of wild-type *C.t.* or CT181::*bla*. On day 7, mice were sacrificed, and the female genital tract was collected in PBS. The tissue was weighed and digested with collagenase/DNAse to release the cells. Cells were lysed using sonication for 10 min to release the bacteria. The lysate was centrifuged at 2,000 x g for 10 min and the supernatant was serially diluted and applied to confluent monolayers of McCoy cells. Inclusions formed after 30 h incubation were counted and plotted.

### Bacterial load determination

Quantitative PCR was performed on homogenized tissue using the QIAamp DNA Mini Kit (Qiagen, CA USA) as described elsewhere (36). Briefly, *C.t.* and mouse DNA was isolated from the tissue. DNA was subjected to quantitative, real-time PCR in triplicate on a StepOne Plus thermal cycler (Applied Biosystems) using the primers listed in Table S1. In parallel, standard curves were generated from known amounts of *C.t.* DNA and mouse DNA and used to determine the amount of DNA present in the sample. Bacterial load was determined by calculating (pg) of *C.t.* DNA per unit weight of (μg) of mouse DNA in the samples.

### Flowcytometry analysis

Mice, infected for 7 days, were sacrificed and the female genital tract was collected in PBS with gentamycin. The tissue was digested with collagenase/DNAse to release the cells and then passed through 70-micron filter to remove undigested tissue. The cell suspension was washed with FACS buffer and stained with CD16/32 (BioLegend Cat#101302) at 1:100 dilution for 30 min on ice. Cells were washed and incubated with CD45-PE594 (BioLegend Cat#103145), CD11b-PEcy7 (ThermoFisher Cat#25-0112-82), F4/80-AF700 (ThermoFisher Cat#56-4801-82) and Ly6G-FITC (BioLegend Cat#127605). CD45^+^ CD11b^+^ Ly6G^+^F4/80^-^ cells were considered as neutrophils.

### ELISA

Cells from the lumen of the female genital tract were collected by flushing with 1 ml of PBS and subsequent centrifugation to pellet down the cells. The secreted cytokines in the supernatant were detected by ELISA. Briefly, IL-10 and IL-6 antibodies were coated on to ELISA plates in coating buffer and incubated overnight at 4°C. The plates were washed three times with PBS/0.05% tween (PBST) and incubated with blocking buffer for 1 h at room temperature. The collected lavage was diluted in the blocking buffer, added to the plates and incubated overnight at 4°C. The plates were washed three times with PBST and incubated for 1 h at room temperature with biotin labelled IL-10 and IL-6 antibodies, respectively according to manufactures protocol. After washing three times with PBST, streptavidin-HRP was added and incubated for 30 min at room temperature. Plates were washed, and the HRP substrate 3,3′,5,5′-Tetramethylbenzidine (TMB) was added. The reaction was stopped by adding stop solution (1N H_2_SO_4_) and the amount of converted TMB was measured at 450 nm. The values obtained were compared to standards to calculate the concentration of cytokines present in the lavages.

## RESULTS

### Identification of secreted effector proteins

Delivery of effector proteins into a host cell during *C.t.* invasion and modulation of early infection events does not require *de novo* protein synthesis, suggesting *C.t.* pre-packages effector proteins at the end of its developmental cycle to initiate new rounds of infection. Indeed TmeA, TmeB, TarP, and TepP are prepacked into EBs at the end of the developmental cycle and have since been confirmed to be secreted proteins required for invasion or promoting the early steps of infection (23, 24, 37, 38). Proteomic profiling of *C.t.* developmental forms identified additional hypothetical proteins that are uniquely present in EBs (Table 1) (25), potentially representing novel effector proteins. To determine whether any of these proteins are delivered into the host cell during *C.t.* infection we conducted CyaA, BlaM, and GSK secretion assays. Expression of the fusion protein was confirmed by western blotting (Fig. S1, S2). CT053 was used as a positive control, which we have previously demonstrated is secreted (35).

**Table 1:** Early effector candidates assayed for secretion. Assay results were not determined (ND) due to ^a^lack of clone, ^b^lack of *C.t.* transformants, ^c^candidate was not expressed, or ^d^candidate was not tested because it was secreted in two prior assays.

| D/UW-3/CX | L2/434/Bu | MW | CyaA Assay | BlaM Assay | GSK Assay | Final Designation |
| --- | --- | --- | --- | --- | --- | --- |
| CT053 | CTL0309 | 17.2kDa | Secreted | Secreted | ND<sup>d</sup> | Secreted |
| CT181 | CTL0433 | 24.4kDa | Secreted | Secreted | ND<sup>d</sup> | Secreted |
| CT365 | CTL0619 | 61.0kDa | ND<sup>c</sup> | ND<sup>c</sup> | Not Secreted | Unable to conclude |
| CT389 | CTL0645 | 47.0kDa | Not Secreted | ND<sup>a</sup> | ND<sup>c</sup> | Unable to conclude |
| CT590 | CTL0853 | 109.1kDa | ND<sup>a</sup> | Not Secreted | Not Secreted | Not Secreted |
| CT668 | CTL0037 | 24.4kDa | Not Secreted | Not Secreted | ND<sup>c</sup> | Not Secreted |
| CT676 | CTL0045 | 19.9kDa | Not Secreted | ND<sup>a</sup> | ND<sup>c</sup> | Unable to conclude |
| CT814 | CTL0185 | 11.4kDa | Not Secreted | Not Secreted | Secreted | Possibly Secreted |
| CT837 | CTL0209 | 76.4kDa | ND<sup>c</sup> | Not Secreted | ND<sup>c</sup> | Unable to conclude |
| CT865 | CTL0244 | 37.5kDa | Not Secreted | Secreted | Not Secreted | Possibly Secreted |

cAMP assays revealed an increase in cAMP for CT181-CyaA relative to CyaA alone, indicating it is released into the host cell during chlamydial infection (Fig. 1A, Table 1). In line with our CyaA assay, our BlaM assay results indicated that CT181-BlaM is secreted (Fig. 1B, Table 1), confirming it represents a *bona fide* secretion substrate. Furthermore, results from our BlaM assay suggests that CT865 may also be secreted. We (28) and others (29, 39) have used the small 13-residue GSK-tag to identify secreted *Chlamydia* proteins. Here we employed this third assay for CT865 as it had conflicting results between the CyaA and BlaM assays. We also used this assay for those that were unable to be assayed or were found to be not secreted using the CyaA and BlaM assays (Table 1). Uniquely, this assay identified CT814 as a potential secretion substrate (Fig. 1C).

**Figure 1:**
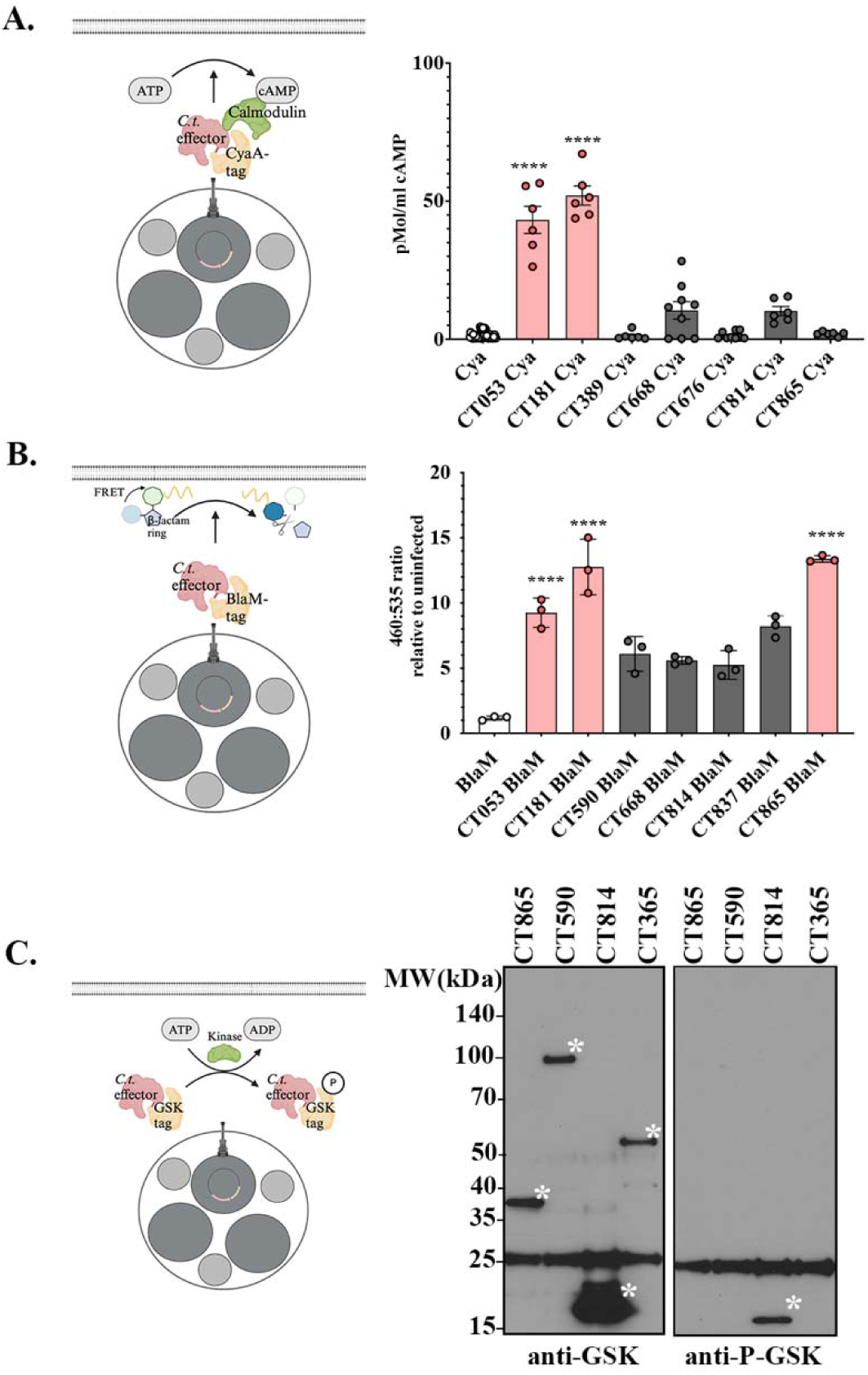
Several proteins, uniquely produced by EBs, are secreted effectors. HeLa cells were infected at an MOI of 5 with each strain for 24h. (**A**) Cytosolic levels of cAMP were measured, and levels obtained from cells infected with the effector-CyaA fusion were compared to cells infected with *C.t.* expressing CyaA alone. Data are from 3 experiments with 3 replicates per experiment. (**B**) Secretion of effector-BlaM fusion was determined by evaluating the change in 460/535nm fluorescence, which results from cleavage of the CCF4-AM substrate. Ratios associated with the effector-BlaM fusion were compared to cells infected with *C.t.* expressing BlaM alone. Data are representative of 3 experiments with 3 replicates per experiment. (**C**) Effector-GSK FLAG fusions were immunoprecipitated using FLAG magnetic beads and analyzed by western blotting using anti-GSK-3β and anti-Phospho-GSK-3β antibodies. Data are representative from three independent experiments. (**A, B**) Error bars represent standard deviation from the means. Statistical significance was determine using One-Way ANOVA comparing the candidate effector to (**A**) CyaA or (**B**) BlaM alone using Dunnett’s multiple comparison post-test. ****P<0.0001.

Collectively, our secretion assays identified CT181 as a new secreted effector that was secreted in two assays. Additionally, we identified two candidates, CT814 and CT865, that were secreted in one assay, and thus might be secreted. Taken together our results highlight the importance of using multiple assays to identify secretion substrates.

### CT181 is important for intracellular replication and inclusion development in cervical epithelial cells

To determine whether the newly identified secretion substrate CT181 is important for chlamydial infection, we used the TargeTron (30, 40, 41) system, successfully generating a CT181 mutant (CT181::*bla*). We then evaluated the ability of the CT181::*bla* mutant to invade, replicate, and form a spacious inclusion relative to wild-type *C.t.,* a CT144 mutant (CT144::*bla*) as an effector control, and a TmeA mutant (*tmeA*-lx) which was used as a positive control for an invasion defect (21, 22, 42). Evaluation of CT181 growth revealed it is important for intracellular replication in A2EN and HeLa cells (Fig. 2A). The growth defect was not simply due to a defect in bacterial invasion, as no significant difference in bacterial uptake was noted for CT181::*bla* compared to wild-type (Fig. 2B). In line with the observed reduction in bacterial replication, smaller inclusions were formed by CT181 mutant relative to wild-type (Fig. 2C, D). Taken together these data indicate that CT181 is important for bacterial replication and inclusion development but is dispensable for host cell invasion.

**Figure 2:**
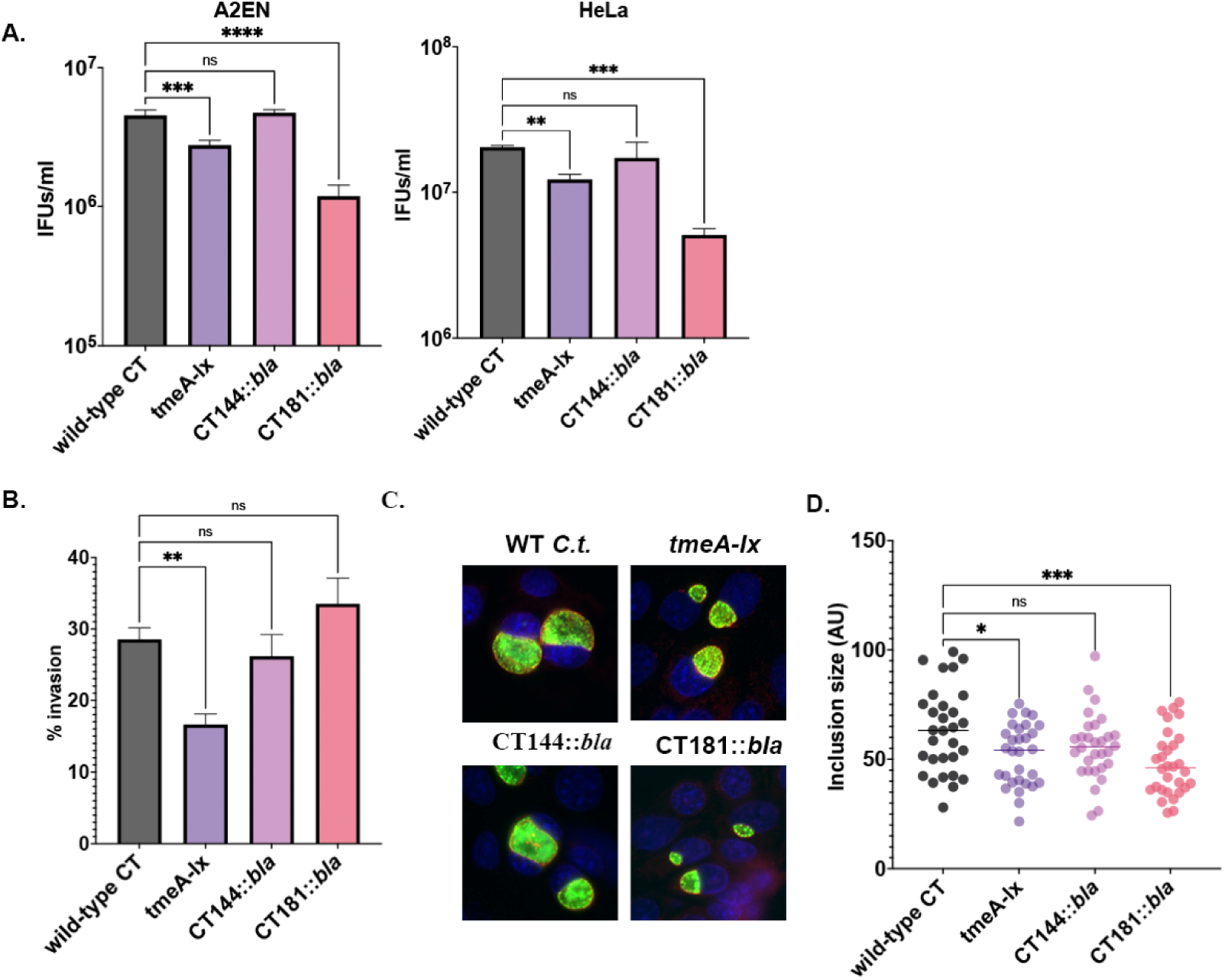
CT181 is dispensable for host cell invasion but is important for intracellular replication and inclusion formation. (**A**) A2EN or HeLa cells were infected at a MOI of 2.5 with wild-type, *tmeA-lx*, CT181::*bla*, or CT144::*bla*. At 48h, cells were lysed and replated on fresh HeLa cell monolayers and infectious forming units were quantified by immunofluorescence microscopy. (**B**) A2EN or HeLa cells were infected at a MOI of 2.5 for 60 min with wild-type, *tmeA-lx*, CT181::*bla*, or CT144::*bla*. The number of internal bacteria was determine using differential immunostaining. (**C)** To measure inclusion size, A2EN cells were infected at an MOI of 2.5 for 24 h. Bacteria were stained with anti-LPS (green), the inclusion membrane was stained with an anti-IncE antibody (red), and DNA was stained with DAPI (blue). Inclusion diameter was measured in ImageJ. (**A-C**) Data are representative of three independent experiments. Statistical significance was determined using One-Way ANOVA with Dunnett’s multiple comparison post-test comparing the mutants to wild-type. ****P<0.0001, ***P<0.001, **P<0.01, *P<0.05.

### CT181 binds to the induced myeloid leukemia cell differentiation protein (Mcl-1)

To dissect the molecular function of CT181, we sought to identify the host target(s). HeLa cells were infected for 24h with *C.t.* L2 expressing FLAG-tagged CT181 under the control of a tetracycline inducible promoter. Affinity purification was performed using FLAG beads and mass spectrometry data were compared to cells infected with *C.t.* harboring the pBomb4-tet vector. A total of 55 proteins were present in all three replicates of CT181, of which 33 were unique to the CT181 AP-MS and were not found in the vector IP (Table S2).

Simultaneous to our AP-MS, we employed a yeast two-hybrid (Y2H) screen to identify potential interacting partners. Several candidates were identified (Table 2), of which the induced myeloid leukemia cell differentiation protein 1 (Mcl-1) represented the largest population of clones and had the highest score. Cross-comparison of hits between the Y2H and AP-MS revealed Mcl-1 as the only hit common to both screens. Interactions between Mcl-1 and CT181 were further supported by alpha-fold modeling (Fig. S3).

**Table 2:** Candidate CT181 binding partners identified by yeast 2-hybrid. Predicted biological score (A-F) was assigned by Hybrigenics. A-Very high confidence in the interaction; B-High confidence in the interaction; C-Good confidence in the interaction; D-Moderate confidence in the interaction; E-Potential non-specific interaction; and F-Experimentally proven artifact.

| Protein | Name | Function | #Clones | PBS Score |
| --- | --- | --- | --- | --- |
| ARMCX6 | armadillo repeat containing X-linked 6 | Regulates mitochondria dynamics and distribution in neural cells | 2 | D |
| BIRC6 | baculoviral IAP repeat containing 6 | Regulation of apoptosis | 5 | D |
| CIR1 | Corepressor interacting with RBPJ | Regulation of transcription | 1 | D |
| DHX29 | DExH-box helicase 29 | Regulation of translation | 3 | D |
| EHBP1 | EH domain-binding protein 1 | Actin cytoskeleton organization | 1 | D |
| ELP1 | Elongator complex protein 1 | tRNA binding | 1 | D |
| FEZ2 | Fasciculation and elongation protein zeta-2 | Signal transduction | 1 | D |
| FOXRED2 | FAD-dependent oxidoreductase domain-containing protein 2 | Ubiquitin-dependent ERAD pathway | 3 | D |
| GPATCH8 | G patch domain-containing protein 8 | RNA binding | 1 | D |
| HNRNPF | Heterogeneous nuclear ribonucleoprotein F | RNA processing | 2 | D |
| LDLRAP1 | Low density lipoprotein receptor adaptor protein 1 | Cholesterol homeostasis | 2 | D |
| LHX9 | LIM/homeobox protein Lhx9 | Regulation of transcription | 1 | D |
| MCL1 | Induced myeloid leukemia cell differentiation protein | Regulation of apoptosis | 81 | A |
| MYCBP2 | E3 ubiquitin-protein ligase | Protein ubiquitination | 7 | C |
| NUMB | Protein numb homolog | Adherens junction organization | 2 | D |
| PGK1 | Phosphoglycerate kinase 1 | Epithelial cell differentiation | 3 | D |
| PRKD3 | Serine/threonine-protein kinase D3 | Protein kinase D signaling | 5 | D |
| PSD3 | PH and SEC7 domain-containing protein | Regulation of ARF protein signal transduction | 2 | D |
| PTBP1 | Polypyrimidine tract-binding protein 1 | RNA splicing | 21 | C |
| SMURF2 | E3 ubiquitin-protein ligase SMURF2 | Protein ubiquitination | 6 | C |
| TMOD3 | Tropomodulin-3 | Actin filament organization | 11 | B |
| WNK1 | Serine/threonine-protein kinase WNK | Ion homeostasis | 1 | D |

To validate this interaction, HeLa cells were co-transfected with GFP-tagged CT181 and HA-tagged Mcl-1. GFP-tagged CT181, or the negative controls GFP-TmeA or GFP alone, were immunoprecipitated from cells and western blots were probed for HA-tagged Mcl-1. GFP-CT181, and not the negative controls, immunoprecipitated HA-Mcl-1 (Fig. 3A) confirming the interaction between these two proteins. To confirm this interaction occurs during infection, HeLa cells were infected at an MOI of 2 with *C.t.* expressing FLAG-tagged CT181, CteG, or TmeA. As shown in Fig. 3A, CT181-FLAG specifically bound to endogenous Mcl-1, confirming their interaction during *C.t.* infection.

**Figure 3:**
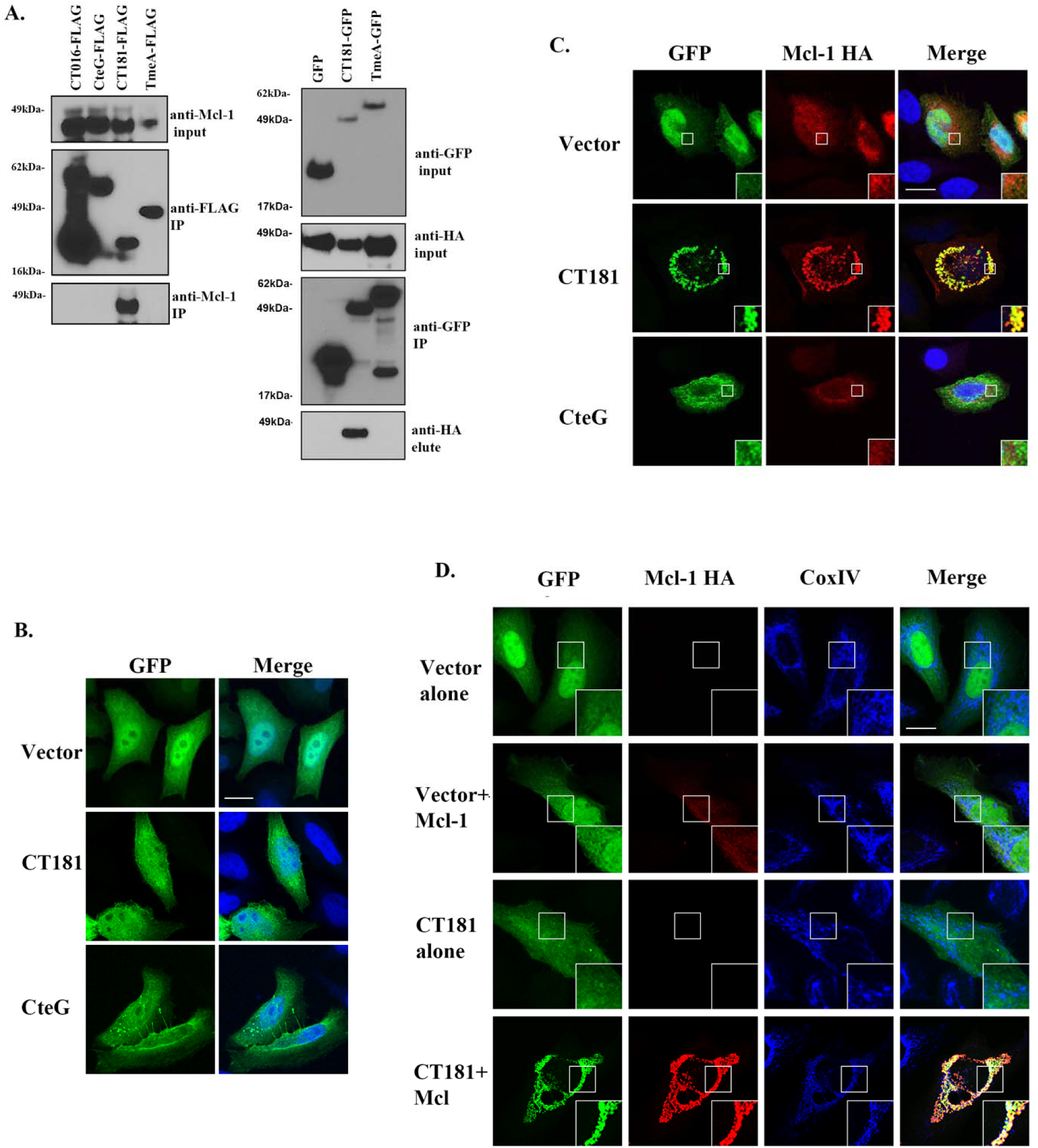
CT181 binds to the induced myeloid leukemia cell differentiation protein. (**A**) HeLa cells were infected with CT016-FLAG, CteG-FLAG, CT181-FLAG, or TmeA-FLAG (left) or co-transfected with HA-tagged Mcl-1 and GFP, GFP-CT181, or GFP-TmeA (right). Proteins were immunoprecipitated using anti-HA beads and analyzed by western blotting using anti-GFP or anti-HA antibodies. Data are representative of three independent experiments. (**B-D**) HeLa cells were transfected with the GFP-effector or GFP-CT181 alone (green) or in combination with HA-tagged Mcl-1. At 24 h post-transfection, cells were fixed with 4% formaldehyde, permeabilized with 0.1% Triton-X 100. (**C**) DAPI (blue) was used to demark the nucleus and Mcl-1 (red) was visualized using anti-HA antibodies. (**D**) Mitochondria were visualized using anti-CoxIV (blue) and Mcl-1 (red) was visualized using anti-HA antibodies. (**B**) Images were obtained using epifluorescence microscopy. Scale bar is 10μm. (**C, D)** Images were obtained using confocal microscopy. Scale bars are 10μm.

### CT181 co-localizes with Mcl-1

To orthogonally confirm that CT181 interacts with Mcl-1 and to gain insight into its potential function, we ectopically expressed GFP-CT181 with or without HA-tagged Mcl-1 and evaluated subcellular localization by immunofluorescence microscopy. When expressed alone, GFP-CT181 was localized throughout the cell (Fig. 3B), however co-expression with Mcl-1 resulted in the formation of large aggregate-like structures (Fig. 3C). Aggregate formation when co-expressed with Mcl-1 was specific to CT181 and was not noted when Mcl-1 was co-expressed with vector or CteG, a *C.t.* effector that binds to CETN2 (34). We also noted that co-expression of Mcl-1 with CT181 increased the Mcl-1 signal (see Fig. 3C, D CT181-Mcl-1 vs. CteG-Mcl-1 or Vec-Mcl-1), suggesting it might stabilize Mcl-1. Mcl-1 is a critical regulator of numerous host cell processes, including apoptosis, mitophagy, mitochondrial bioenergetics, cell cycle, and DNA repair (43). To gain insight into the role Mcl-1 might play during chlamydial infection, we co-expressed CT181 with Mcl-1 and assessed co-localization with the mitochondrial marker CoxIV. While CT181 did not co-localize with CoxIV when expressed independently of Mcl-1, co-expression with Mcl-1 resulted in prominent co-localization with CoxVI (Fig. 3D). Again, we noted significantly increased signal intensity when Mcl-1 was co-expressed with CT181 but not vector (Fig. 3D). Collectively, these results indicate that CT181 binds to Mcl-1 and may act to stabilize Mcl-1.

### CT181 is involved in PMN lifespan extension

Previous work has shown that *Chlamydia* infection can interfere with host cell death induced by extrinsic or intrinsic stress stimuli (13–15, 44). To investigate the role of CT181 in regulation of infection-induced cell death, we first tested whether CT181 could protect cells from apoptotic stimuli or whether CT181 was necessary for *C.t.* resistance to apoptotic stimuli. Surprisingly, ectopic expression of CT181 was unable to protect cells from apoptotic stimuli and no difference in sensitivity for CT181::*bla* mutant relative to wild-type *C.t.* was noted following apoptotic stimulation with staurosporine (data not shown). Unlike epithelial cells, polymorphonuclear neutrophils (PMNs) have a short lifespan of only 12 to 24 hours in the bloodstream without stimulation and are inherently programmed to undergo spontaneous apoptosis (45). Notably, Mcl-1 plays a central role in PMN survival (45). We have previously shown that *C.t.* can survive in PMNs for extended periods, even beyond 24 h (46), suggesting that infection may prolong PMN lifespan. We therefore investigated whether CT181 extends the lifespan of human PMNs. Human PMNs were infected with wild-type *C.t.* or CT181::*bla* and cell viability was analyzed by flow cytometry (Fig. 4A). Wild-type *C.t.*, but not CT181*::bla*, significantly extended the survival of PMNs, with more than 50% of the cells remaining alive even after 48 hours (Fig. 4B). Next, we investigated whether infection of human PMNs affects the amount of Mcl-1. Due to the highly variable nature of PMNs, we infected PMNs isolated from the blood of different donors. A strong increase in Mcl-1 levels was detected upon wild-type *C.t.* infection. Interestingly, CT181::*bla* infection failed to increase Mcl-1 levels in PMNs from three different donors (Fig. 4C-D). These results are consistent with a strongly reduced survival of the CT181::*bla* in human PMNs, with only 35 % of the initial infection inoculum being detected at 48 hpi and no detectable infectious bacteria at 96 hpi (Fig. 4E-I). Taken together, our results suggest that CT181 is important for *C.t.*’s ability to prolong survival of human PMNs, possibly by stabilizing Mcl-1.

**Figure 4:**
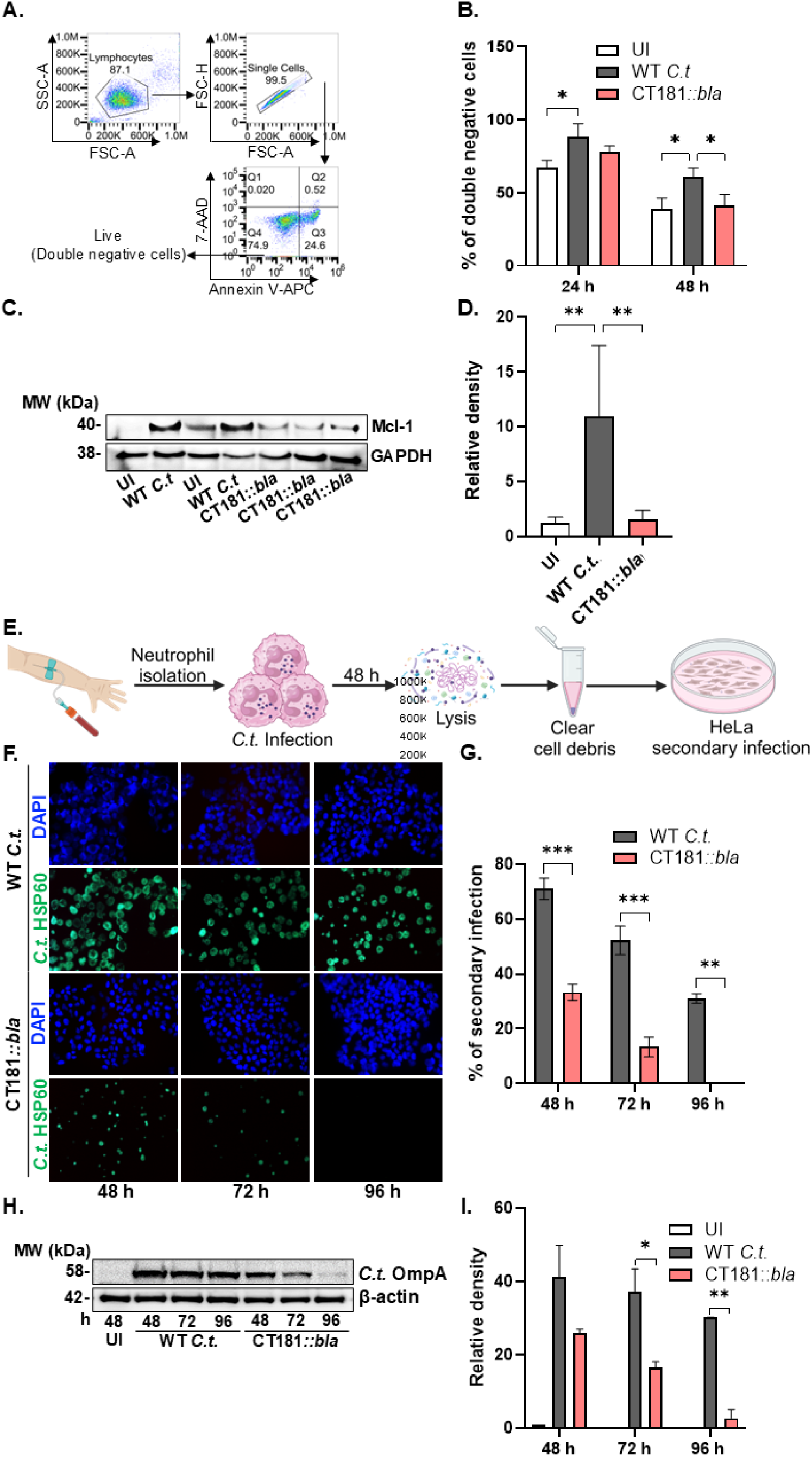
CT181 is involved in the prolongation of human PMN lifespan and the survival of *C.t*. Freshly isolated PMNs from human blood were infected with wild-type (WT) *C.t.* or CT181::*bla* at an MOI of 5 for the indicated time points. (**A**) PMNs were stained with Annexin V and 7-AAD at the indicated time points and analyzed for cell survival using flow cytometry. **(B)** Annexin V- and 7AAD-negative cells were considered as live cells and shown as percentage of double negative cells. (**C**) Mcl-1 expression was analyzed by western blot 48 hours post infection (h.p.i). The data are representative of three independent experiments. **(D)** Relative densities were quantified and plotted using ImageJ. **(E)** Graphical representation of the work plan for re-infectivity assays. Freshly isolated human PMNs were infected with *C.t.* or CT181::*bla* for 48 h, lysed and released bacteria were added to fresh HeLa cells. (**F**) Infected PMNs were lysed and replated on fresh HeLa cell monolayers and infectious forming units were quantified using immunofluorescence. Bacteria were stained with anti-OmpA (green) and nuclei with DAPI (blue). (**G**) Inclusion counts were calculated and plotted as percent of infection. (**H**) In a replicate experiment, HeLa cells were harvested to determine the total *C.t.* OmpA expression relative to human Actin levels to estimate infectious burden. The data are representative of two independent experiments. Statistical significance was determined using Two-Way ANOVA with Tukey’s multiple comparison test. ****P<0.0001, *P<0.05.

### CT181 is required for survival of *C.t.* in mouse PMNs and in transcervical infection

To investigate the role of CT181 in the interaction of *C.t.* with PMNs in an immunocompetent model, we repeated experiments performed with human PMNs using mouse PMNs. Similar to human PMNs, infection with wild-type bacteria extended the lifespan of mouse PMNs at 48 hpi but had no effect at 24 hpi (Fig. 5A). The CT181::*bla* mutant failed to extend the lifespan of mouse PMNs at any of the investigated time points and the number of viable cells was lower than that of the non-infected PMN sample at 24 hpi (Fig. 5A). Fewer neutrophils survived during CT181::*bla* infection, possibly because the absence of CT181 failed to stabilize Mcl-1, as evidenced by decreased Mcl-1 expression in CT181::*bla*-infected neutrophils compared to wild-type-infected neutrophils (Fig. 5B-C). CT181 was important for *C.t.* survival in PMNs as significantly fewer mutant bacteria survived exposure to mouse PMNs compared to wild-type *C.t* (Fig. 5D-H). In addition, we infected mice by transcervical infection and determined the number of infectious EBs by infecting McCoy cells with lysates of the female genital tract (FGT) 6 days pi (Fig. 6A). Significantly fewer infectious EBs could be isolated from mice infected with the CT181::*bla* mutant compared to those infected with the wild-type bacteria (Fig. 6B-C). Interestingly, the number of PMNs detected in the FGT tissue of the mice infected with CT181::*bla* was significantly lower compared to the wild-type infected mice (Fig. 6D-E). The lower number of PMNs correlated with lower levels of the inflammatory cytokines TNFα and IL-6 (Fig. 6F-G) suggesting that CT181::*bla* is severely affected in the PMN response.

**Figure 5:**
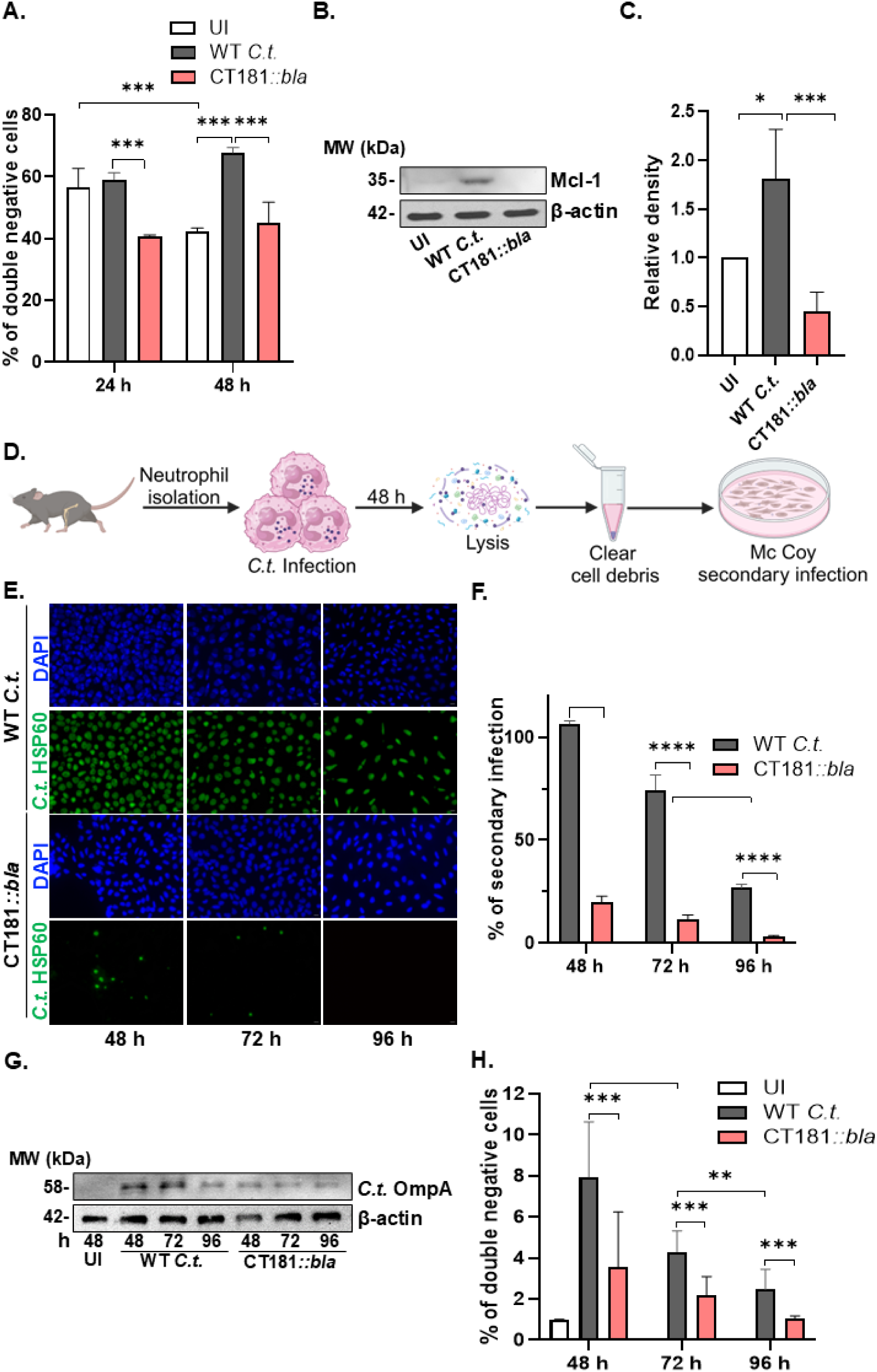
CT181 is involved in the prolongation of mouse PMN lifespan and the survival of *C.t.*. Neutrophils were isolated from the bone marrow of naïve C57BL6/N mice and were infected with wild-type or CT181::*bla C.t.* at an MOI of 5 for the indicated time points. (**A**) PMNs were stained with Annexin V and 7-AAD and analyzed for cell survival using flow cytometry. Double negative cells were considered as live cells and shown as percentage of double negative cells. (**B**) Mcl-1 expression was analyzed by western blot at 48 hpi. The data are representative of three independent experiments. **(C)** Relative densities were quantified and plotted using ImageJ. **(D)** Graphical representation of the work plan for re-infectivity assays. Freshly isolated human PMNs were infected with *C.t.* or CT181::*bla* for 48 h, lysed and released bacteria were added to fresh McCoy cells. (**E**) Infected PMNs were lysed and replated on fresh McCoy cell monolayers, and infectious forming units were quantified by immunofluorescence. Bacteria were stained with anti-OmpA (green) and nuclei with DAPI (blue). (**F**) Inclusion counts were calculated in ImageJ and plotted as percentage of infection. (**G**) In a replicate experiment, McCoy cells were harvested to determine the total *C.t.* OmpA expression relative to mouse actin levels to estimate infectious load. The data are representative of three independent experiments. **(H)** Relative densities were quantified and plotted using ImageJ. Statistical significance was determined using Two-Way ANOVA with Tukey’s multiple comparison test. ****P<0.0001, ***P<0.001, **P<0.01, *P<0.05.

**Figure 6:**
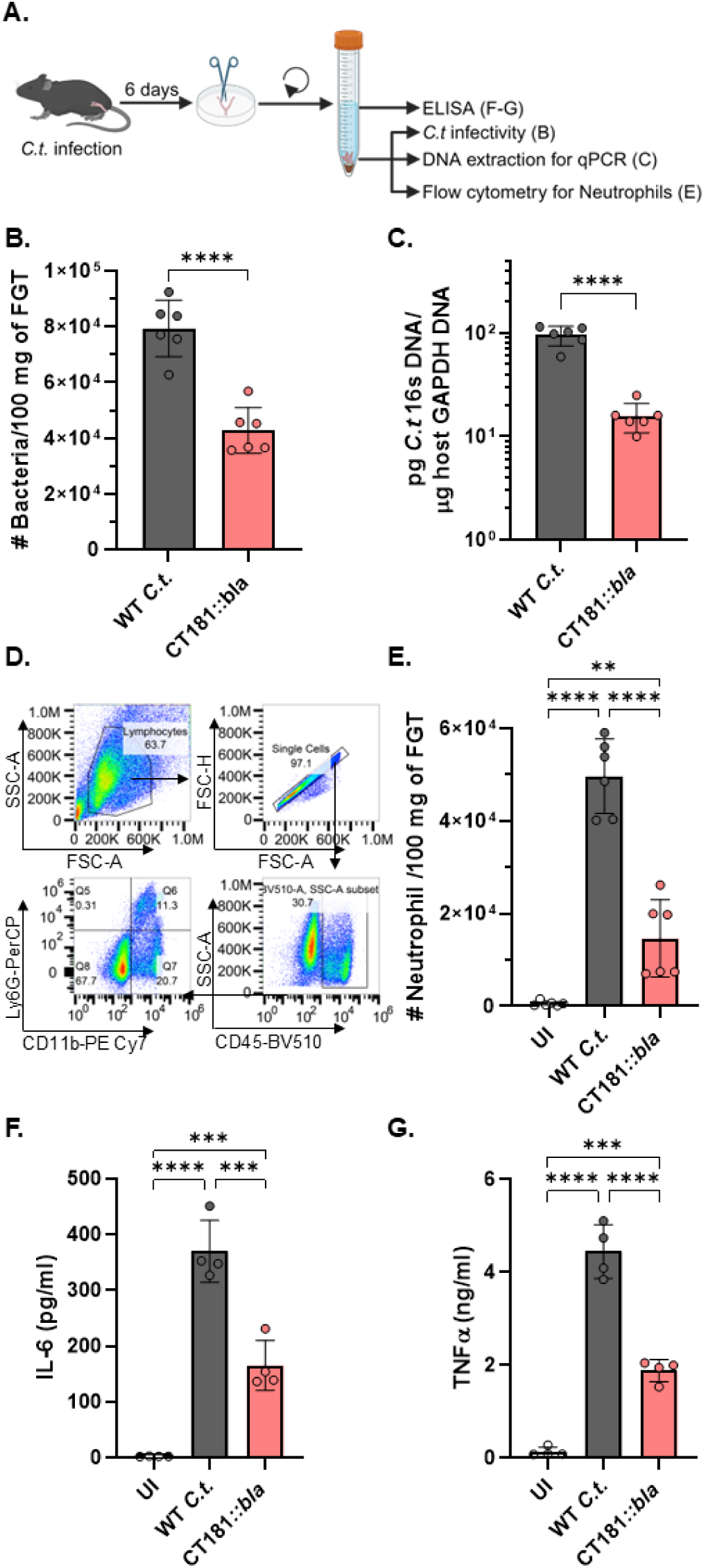
CT181 is involved in survival of *Chlamydia* during mouse genital tract infection. Female C57BL/6N mice were infected for 6 days with 10^7^ wild-type or CT181::*bla C.t*. (**A**) Graphical representation of the experimental setup. At 6dpi, the female genital tract was harvested, weighed, and lysed using collagenase D/DNase I. (**B**) A portion of the cells were collected and lysed by sonication. The lysate and released *C.t*. were added to a monolayer of McCoy cells. The number of inclusions was counted from each mouse and expressed as the number of bacteria per 100 mg of tissue. (**C**) A portion of the tissue was subjected to DNA isolation, and the bacterial load was determined through quantitative real-time PCR of the *C.t.* 16s RNA gene and the mouse GAPDH gene. (pg) *C.t.* 16s RNA gene / ug host GAPDH gene was plotted to determine bacterial burden in the tissue. (**D**) A portion of the harvested cells were stained for CD45, CD11b, Ly6G, CD62L, and F4/80. CD45+ CD11b+ F4/80-Ly6G+ CD62L+ were considered as neutrophils recruited to the tissue. Shown is the gating strategy for the neutrophil count. (**E**) The total number of cells collected was analyzed and expressed as the number of neutrophils per 100 mg of tissue. The data points represent the results obtained from a single mouse (n=6). (**F**) IL-6 and (**G**) TNFα levels were determined from female genital tract lavages by ELISA (n=4). Statistical significance was determined using One-Way ANOVA with Tukey’s multiple comparison test. ***P<0.001, **P<0.01.

## DISCUSSION

Infection of a host cell by an obligate intracellular pathogen requires the pathogen to subvert host cell death pathways while simultaneously promoting host cell viability, a phenotype conferred by bacterial effector proteins. Although Inc proteins have been readily identified based on the presence of a bi-lobed hydrophobic domain, the identification of non-Inc secreted factors has proven more challenging. By focusing on proteins uniquely present in the EB of *C.t.*, we identified CT181 as a novel secreted effector protein and demonstrate that it binds to Mcl-1. Our findings reveal a previously unknown mechanism whereby CT181 prolongs neutrophil survival through stabilization of Mcl-1. Furthermore, we show that infection with the CT181 mutant leads to a significant reduction in the number of PMNs and a decrease in the production of inflammatory cytokines, implicating CT181 in modulating the host immune response to *C.t.* infection and underscoring its dual role in promoting bacterial survival and influencing the host’s inflammatory environment.

To date, only a handful of secreted effectors have been identified and characterized, limiting our understanding of the roles they play in *C.t.* pathogenesis. While some effectors have been identified based on the presence of a eukaryotic-like domain or via their interaction with chaperones (17–19, 24, 47, 48), it is likely that other *C.t.* secreted proteins remain to be discovered. Prior work has demonstrated that effector proteins associated with host cell invasion and early infection events are uniquely prepackaged at the end of the developmental cycle, positioning them to initiate new rounds of infection (23, 24, 42). While TarP and TmeA directly promote host cell invasion by inducing cytoskeletal rearrangements (21–23, 42, 49–52), TmeB appears to play an opposing role by interfering with Arp2/3-mediated actin polymerization (53). TepP represents another multifunctional prepackaged effector that binds to Crk adaptor proteins and the PI3K complex, stimulating PI(3,4,5)P_3_ formation, and it disassembles tight junctions by perturbing EPS8 (24, 38, 54). Intriguingly, TepP also regulates the expression of immunity-regulated genes and reduces neutrophil recruitment in an organoid infection model (24, 55). Given the diverse roles these early effector proteins play in promoting *C.t.* infection, we rationalized additional hypothetical proteins uniquely produced by EBs may represent novel secreted effector proteins (25). By screening 10 candidate proteins for secretion, we identified CT181 as a novel secretion substrate. Our data indicate that CT181 is dispensable for invasion but might be important later in infection.

Manipulation of host cell viability is a critical defense mechanism employed by a broad range of microbial pathogens. While some pathogenic bacteria, such as *Yersinia*, *Shigella,* and *Salmonella,* promote apoptosis to induce killing of phagocytes, obligate intracellular bacteria like *C.t.* have evolved sophisticated strategies to protect their host cells from apoptotic stimuli while simultaneously stimulating pro-survival pathways. During apoptosis, pro-apoptotic Bcl-2 family members Bax and Bak trigger permeabilization of the outer mitochondrial membrane, a critical step in the apoptotic cascade. However, anti-apoptotic Bcl-2 family members, including Mcl-1, counteract this process by sequestering Bax and Bak in the cytoplasm (43). Mcl-1’s expression and activity are tightly controlled at multiple levels - transcriptional, posttranscriptional, and translational - allowing for finely tuned and rapid changes in response to internal and external stimuli (43). Crucially, Mcl-1 is further regulated via rapid protein turnover through ubiquitination and proteasomal degradation, making it an ideal target for pathogens seeking to manipulate host cell survival. During *C.t.* infection, both mRNA and protein levels of Mcl-1 are up-regulated in a RAF/MEK/ERK-pathway-dependent manner (15). Activation of this signaling pathway results in Mcl-1 phosphorylation and stabilization, enhancing its anti-apoptotic activity and ensuring survival of the host cell. Interestingly, Mcl-1 is also stabilized during *C.t.* infection via deubiquitination at the inclusion membrane by the *Chlamydia* deubiquitinating enzyme 1 (Cdu1) (16). While a Cdu1 mutant reduced Mcl-1 levels during infection, Mcl-1 levels remained elevated relative to uninfected cells or those treated with an apoptosis inducer (16). This suggests that *C.t.* employs multiple, potentially redundant methods to maintain Mcl-1 levels. Our study builds upon these findings by demonstrating that the secreted protein CT181 binds to Mcl-1 and that CT181 expression is important for stabilizing Mcl-1 expression. This discovery adds another layer of complexity to *C.t.*-host interactions, revealing a novel mechanism for the pathogen to manipulate host cell survival pathways. The multi-faceted approach taken by *C.t.* in manipulating Mcl-1 levels - through signaling pathway activation, deubiquitylation, and now direct protein-protein interaction - underscores the central role of Mcl-1 in bacterial survival.

Neutrophils are essential components of the innate immune system and as such, play a crucial role in the initial host response to microbial pathogens. These cells are among the first to respond to microbial infection where they act to limit bacterial spread and facilitate pathogen clearance via phagocytosis, degranulation, and release of neutrophil extracellular traps (NETs). Notably, neutrophils have a short lifespan, typically undergoing apoptosis within 12-24 h, at which point scavenger macrophages clear out the apoptotic neutrophils, leading to resolution of the inflammatory response and containment of cytotoxic materials that could otherwise damage host tissues. However, under certain conditions, including during inflammation or infection, the lifespan of neutrophils can be prolonged. *Anaplasma phagocytophilum*, the causative agent of human granulocytic anaplasmosis, induces transcription upregulation of Mcl-1 (56, 57), activates the PI3K/Akt pathway to maintain Mcl-1 expression (58), and induces phosphorylation of p38 mitogen-activated protein kinase (MAPK) (59), all to promote neutrophil survival. *Coxiella burnetii* also prolongs neutrophil survival by activating MAPK pathways to promote stabilization of Mcl-1. While the specific factor involved remains unknown, Mcl-1 stabilization and inhibition of neutrophil apoptosis occur in a type IV secretion system dependent manner, suggesting that a secreted protein is required (60, 61). Additionally, *C. pneumoniae* and *C.t.* can prolong the life span of neutrophils by delaying apoptosis (44, 62, 63). Previous studies have shown that *C.t.* can remain viable and infectious for at least 24 h after infection (46, 64), suggesting prolonging neutrophil survival may serve as a protected niche for the pathogen to spread during infection (9). Remarkably, and consistent with this notion, we have now shown that wild-type *C.t.* can survive in the presence of neutrophils for up to 96 h. The CT181 mutant showed no detectable infectious particles at this time point, which can be attributed to both a failure to prolong neutrophil lifespan and a reduction in *C.t* survival. Collectively, while prior studies highlight the importance of neutrophil survival, the mechanism remained unknown. Our study indicates that secreted bacterial proteins, namely CT181, may be crucial to this process.

In conclusion, we have identified a novel *C.t.* secreted protein, CT181, that binds to Mcl-1 and prolongs neutrophil survival during *C.t.* infection. Our work provides a framework for future studies aimed at understanding how bacterial effectors proteins disarm the host immune response to allow for productive infection.

## Acknowledgments

We acknowledge grant support from the NIH (M.M.W., R01 AI150812, R01 AI155434, and R61 AI179999, B.S. T32 AI007511), the University of Iowa Stead Family Scholars to M.M.W, and the European Research Council (ERC) under the European Union’s Horizon 2020 research and innovation program (ERC-2018-ADG/NCI-CAD) to T.R. We thank Weber lab members Steve Huang, Alix McCullough, Jaby, and Xavier Tijerina for assistance. We thank Kristin Patrick for helpful discussions.

