## Supplemental for "The *Chlamydia trachomatis* secreted effector protein CT181 binds to Mcl-1 to prolong neutrophil survival"

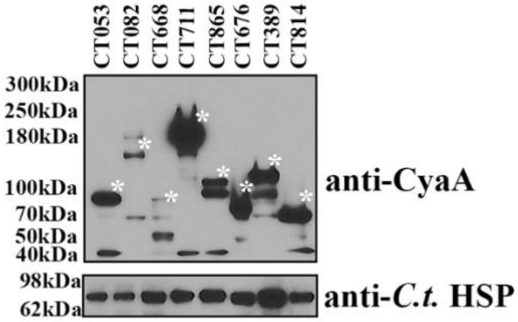


**Figure S1: Expression of candidate effector-CyaA fusion proteins in *C.t.*** Candidate secretion substrates were expressed as C-terminal fusions to the CyaA-tag and transformed into *C.t*. HeLa cells were infected at an MOI of 5 for 24h with each candidate, after which expression of the fusion protein was confirmed by immunoblotting with anti-CyaA antibodies. *C.t.* HSP-60 was used as a loading control. Asterisks on the right of each band indicate the product corresponding to the anticipated molecular weight of each CyaA fusion protein. Data are representative of three independent experiments.

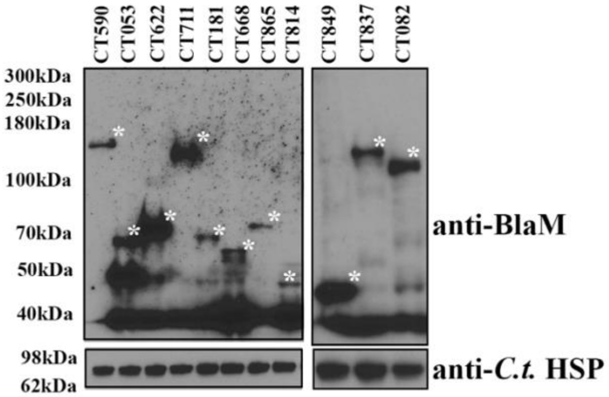


**Figure S2: Expression of candidate effector-BlaM fusion proteins in *C.t.*** Candidate secretion substrates were expressed as C-terminal fusions to the BlaM-tag and transformed into *C.t*. HeLa cells were infected at an MOI of 5 for 24h with each candidate, after which expression of the fusion protein was confirmed by immunoblotting with anti-BlaM antibodies. *C.t.* HSP-60 was used as a loading control. Asterisks on the right of each band indicate the product corresponding to the anticipated molecular weight of each BlaM fusion protein. Data are representative of three independent experiments.

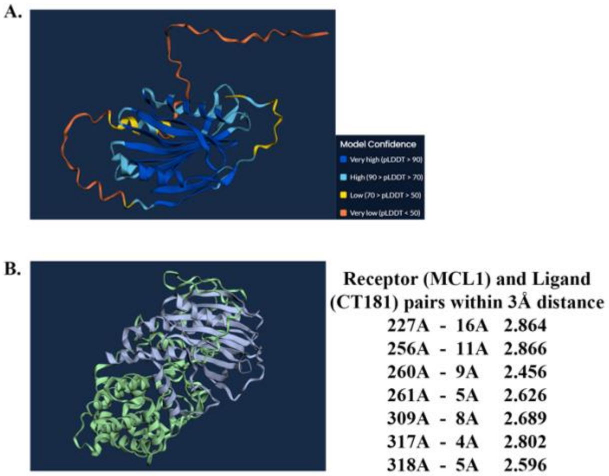


**Figure S3: Alphafold model of CT181 with and without Mcl-1.** Alphafold predictions of CT181 alone or with Mcl-1 (B).
